# Fluid Menisci and *In Vitro* Particle Dosimetry of Submerged Cells

**DOI:** 10.1101/2021.03.25.436962

**Authors:** Sandor Balog, Barbara Rothen-Rutishauser, Alke Petri-Fink

## Abstract

Understanding the mechanisms of interaction between cells and particulate nanomaterials lies in the heart of assessing the hazard associated with nanoparticles. The paradigm of toxicology requires quantifying and interpreting dose-response relationships, and cells cultured *in vitro* and exposed to particle dispersions rely on mathematical models that estimate the received nanoparticle dose. Yet, none of these models acknowledges the fact that aqueous cell-culture media wet the inner surface of hydrophilic open wells, which results in curved fluid-air interface called meniscus. We show that omitting this phenomenon leads to a nontrivial but systematic error and twists the fundamental concept of nanotoxicology. Given that reproducibility and harmonization between meta analyses, *in vitro*, *in silico*, and *in vivo* studies must be improved, we present an adequate mathematical model that greatly advances such efforts.

## INTRODUCTION

*In vitro* assessment of the hazard of nanomaterials^1, 2^ requires predictive particle dosimetry models dedicated to estimate the dose from nanoparticles in dispersion exposure.^3–13^ Let it be either the pharmacologicy^14–17^ or the toxicology of nanomaterials,^18–23^ it is a fundamental theorem that biological response is dose-dependent, and therefore accuracy is most important to *in vitro* exposures.^24–26^ To culture and expose cells, open plastic wells are used whose inner surface is made hydrophilic. Accordingly, cell-culture media wet the inner surface of the wells, which results in a concave fluid surface called meniscus (Figure 1).

**Figure 1.**
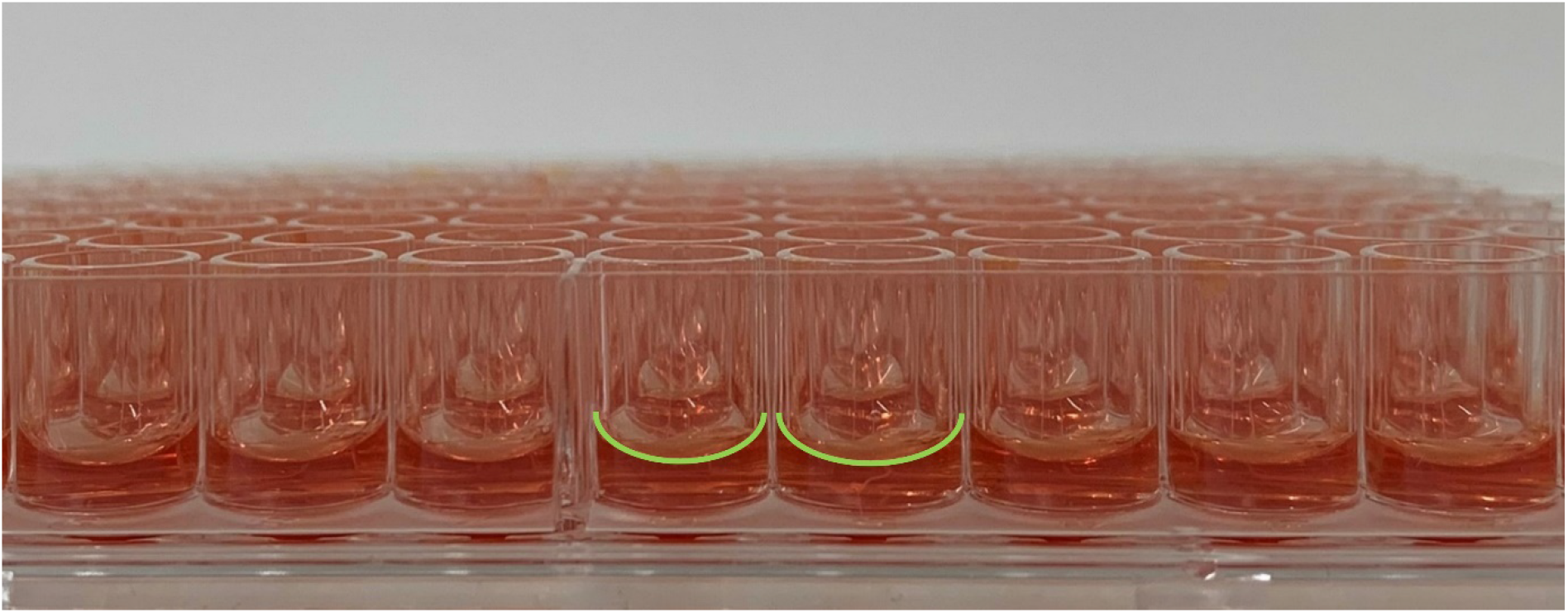
Equilibrium fluid menisci in open polystyrene microplate wells filled with aqueous cell culture medium. Owing to surface tension and wetting phenomena, a concave air-fluid interface, called meniscus, forms spontaneously. The contour of such interface is highlighted in green in two wells in the middle. The level of the fluid in the wells is a function of the distance from the center, and is highest at the wall and smallest at the center. When the wells are cylindrical, the meniscus is axially symmetric with a circular contour.

While this is a fundamental phenomenon that concerns any open well that exhibits a fluid-air interface, mathematical models assume flat fluid surfaces, neglecting completely the meniscus.^3–13^

We will address this deficiency here. First, we assemble the mathematical model—which is a continuous and deterministic model that comprises two second-order nonlinear and linear partial differential equations with corresponding boundary and initial conditions. Our model addresses adequately the meniscus, medium geometry, and the transport equation in three plus one dimensions: spatial coordinates and time. Next, the model is evaluated via finite element method, using characteristic particle sizes and mass densities at systematically varied affinities for particle internalization. Finally, the corresponding solutions obtained by simulations— providing the concentration of particles as a function time and space—are analyzed and the impact of meniscus on *in vitro* particle dosimetry is discussed.

All symbolic and numerical computation is performed by using Mathematica Version 12.0 (Wolfram Language, Wolfram Research, Inc., Champaign, IL).

## MATHEMATICAL MODEL

### Equilibrium shape of the meniscus

First, we obtain the equilibrium shape of the meniscus, and then set up the corresponding geometry we need for solving the particle transport equation with the initial and boundary conditions. Given the scope of our work, we model stable systems of uniform spherical particles that do not disintegrate *e.g.*, via dissolution, and do not aggregate, thus their hydrodynamic properties remain constant. However, it is not difficult to adapt the approach to either non-stationary or anisotropic or polydisperse partcles.^6, 11^ Furthermore, while we dominantly address a quiescent system where there is no collective fluid motion (*e.g.*, convection), we shall in some detail discuss the of potential impact of pipetting. By the term ‘pipetting’ we refer to administration of the particles to the well, which—to our best knowledge—has never been addressed before in the context of *in vitro* particle dosimetry. Finally, we compare the result given by the usual one-dimensional model and the results given by our model developed here. This is especially relevant in view of the fact that one-dimensional models—using a volume-equivalent fluid height—concern hundreds of already published work.

The equilibrium shape of the meniscus may be obtained by solving a nonlinear second-order ordinary differential equation obtained via combining the principle of minimum energy and the Young–Laplace equation (Supporting Information, Equilibrium shape of the meniscus). Here we address cylindrical wells, however, numerical solutions for rectangular shapes may be also readily found. Nevertheless, a well with axial symmetry enables us to use cylindrical coordinates instead of Cartesian coordinates (Figure 2), and given the dimensions of commonly used wells, we may safely assume that the Gaussian curvature of the surface is small (Supporting Information, Dimensions and the working volumes of wells with axial symmetry and flat bottom, Table SI-1). The solution is then

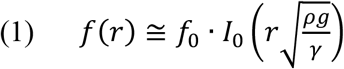

where *I*_*n*_ is the modified Bessel function of the first kind with *n* = 0, *ρ* the mass density of the fluid, and *γ* surface tension. For a volume of *V* filling the well, we find the value of *f*_0_ = *f* (*r* = 0) via the following expression:

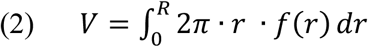

where *R* is the inner radius of the well. After performing the integration on Equation 1, we obtain

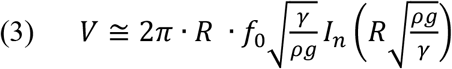

where *I*_*n*_ is the modified Bessel function of the first kind with *n* = 1. We express the value of *f*_0_ via Equation 3, and Figure 2 shows the shape of the meniscus obtained via Equation 1 at 37 °C in a cylindrical well with a radius of 5.5 mm and a fluid volume of 250 μL. For the one-dimensional model, the volume-equivalent fluid height is *ℎ* = *V/*(*π* ∙ *R*^2^), which is the ratio of the fluid volume and the surface area of the well. Surface tension **γ** is a function of temperature, and in the case of an aqueous cell culture medium of 37 °C, *γ* ≅ 7 ∙ 10^−4^ J/m^2^. It is worth mentioning that cell culture media are usually supplemented, *e.g.*, with fetal bovine serum, to promote the growth and maintenance of cells. Supplemented cell culture media contain proteins (*e.g.*, bovine serum albumin), which exhibit both hydrophobic and hydrophilic regions. Such proteins, alike to surfactants, reduce the surface tension at the air-liquid interface, (**γ**) and improve the wetting of the wall.^27^ This aspect must be considered if protein concentration is high.

**Figure 2.**
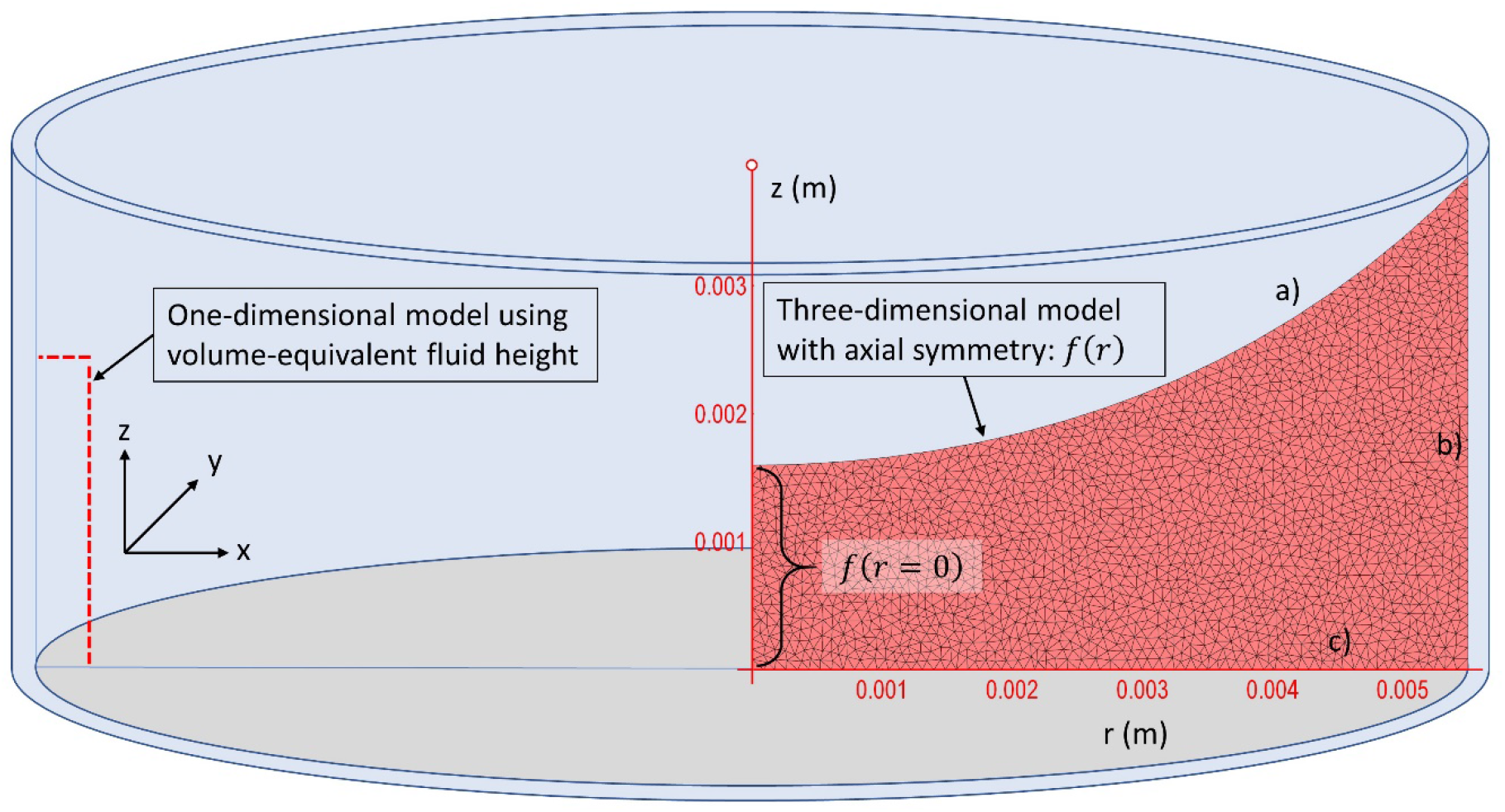
Schematic view of a cylindrical well filled with cell culture medium. The shape of the meniscus shown here is to scale and obtained by using the dimensions of a 48-well plate with a well diameter of 11 mm, and a filling-volume of 250 μL at a temperature of 37 °C. The corresponding volume-equivalent fluid height is 2.63 mm, which value is used by the one-dimensional model (Equation 4) neglecting the phenomenon of fluid meniscus. In an open cell culture well, there are three interfaces: a) fluid-air interface on the top, b) fluid-wall interface, and c) fluid-cell interface on the bottom. The axial symmetry of the three-dimensional geometry enables to use cylindrical coordinates [*r, z*] instead of Cartesian coordinates [*x, y, z*], reducing the dimensionality of the problem, and thus it becomes computationally less expensive. In order to solve the three-dimensional transport equation in this geometry (Equation SI 10 and Equation 5), we use a finite element method, which is particularly useful for solving numerically partial differential equations on nontrivial geometries where a closed-form solution is either unavailable or impractical. Finite element methods approximate continuous functions in space and time with a set of corresponding quantities calculated on discrete points of a grid, such as the triangular-mesh shown in the figure. In case of rectangular wells, continuous rotational symmetry is not present, and one must use Cartesian coordinates.

### Transport equation, initial and boundary conditions

Next, we present the transport equation along with the initial and boundary conditions we use. The transport equation is commonly used in a simple one-dimensional form,^3–13^ using the volume-equivalent fluid height (Figure 2)

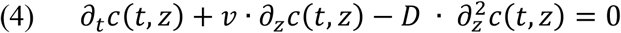

where our work *c*(*t*,…) will represent mass concentration. The three-dimensional model in cylindrical geometry is expressed as (Supporting Information, Transport equation)

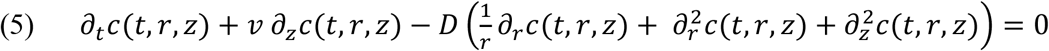

where *D* is the translational self-diffusion coefficient and *υ* is the settling velocity of the particles (Supporting Information, Transport equation).

Before solving Equation 4 and 5, we need to define a) the initial condition, *i.e.*, the value of *c* (mass concentration) throughout the fluid volume at *t* = 0, and b) the boundary conditions at the interfaces. As initial condition, we take that the concentration is identical throughout the fluid volume and the dispersion is quiescent at *t* = 0. Accordingly, we do not address the potential impact of particle administration (namely, pipetting) and its possible impact on particle transport (Supporting Information, Initial condition and the impact of pipetting).

Regarding the boundary conditions, there are three interfaces we need to address: a) fluid-air interface on the top, b) fluid-wall interface on the side, and c) fluid-cell interface on the bottom (Figure 2). The particle cannot cross the fluid-air interface *f*(*r*) on the top, and thus, they bounce back from it. The same applies at the fluid-wall interface on the side if there are no cells adhering to the walls.^28^ This property along *f*(*r*) and along the walls *r* = *R* is expressed by the so-called reflective Neuman boundary condition: *∂_r_c* = 0. The fluid-cell interface is taken as a flat surface at *z* = 0, however, the cells adhering to the bottom are of finite thickness and may also vary in height. Yet, the fluid height (on the order of a few mm) is usually at least three orders of magnitude larger than the height of a cell (on the order of a few μm), and therefore, the error is insignificant. To describe the boundary condition at the fluid-cell interface— depending on *e.g.*, the expected ability and total capacity of the cells to internalize particles— there are several models that have been used previously.^3–13^ The early models assumed that cells internalize particles instantaneously, and the interface is treated as a perfectly absorbing surface using the Dirichlet boundary where the concentration is zero at all times: *c* = 0 at *z* = 0. Later, computational models assumed Neuman boundary condition, also known as flux-type boundary conditions. One may also use a linear combination of the Dirichlet and Neumann boundary conditions, which is referred to as Robin boundary condition. If cells are grown on the wall, the boundary condition is to be expressed similarly as used at the fluid-cell interface on the bottom. Furthermore, to address cellular dose, one might need to consider appropriate boundary conditions for describing the phenomena of particle desorption and exocytosis,^29 30^ *i.e.*, particles adhered to the cell may be released before internalization occurs, and internalized particles may later be secreted by the cell.

Here we will use a Neuman boundary condition: *∂_z_c*(*t, r, z*) < 0. This boundary condition in fact expresses a concentration difference between *c*(*t, r, z* + *dz*) and *c*(*t, r, z*) at *z* = 0 in the limit of vanishing *dz*, and the inequality means that *c*(*t, r, z* + *dz*) < *c*(*t, r, z*), *i.e.*, the cells internalize particles. The Neuman boundary may be used to represent either a constant rate of particle internalization—independently on the concentration right above the cell—or a concentration-dependent rate of internalization, where the dependence may be either linear or nonlinear. To describe this relationship, frequently an expression mimicking the Hill-Langmuir model is used, expressing an equilibrium between adsorptions and desorption in ligand binding.^7, 12^ Here we use the model where the rate of particle internalization is proportional to the mass concentration of particles at the cell:

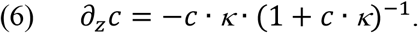

The unit of *κ* > 0 is *m*^−1^, and the larger *κ*, the higher the rate of particle uptake. It is worth mentioning that the Dirichlet condition with *c*(*t, r*, 0) = 0 represents a perfectly absorbing surface and defines the upper limit of the rate of internalization of particles. Accordingly, as *κ* increases, the rate of uptake approaches asymptotically that of a perfectly and instantaneously absorbing surface. We adopt an effective surface coverage equal to one, which means that the bottom of the well is completely and homogeneously covered by cells. This however may be changed to describe partial coverage and heterogeneous cells population in terms of affinity for particles internalization, and the boundary conditions may be formulated accordingly.

Regarding dose of *in vitro* exposure of nanoparticles, one distinguishes administered dose, delivered dose, and cellular dose. The administered dose is proportional to the particle concentration and the volume of the dispersion added to the cell culture well. The delivered dose is proportional to the administered dose and the rate of transport of particles to the cells. The cellular dose is proportional to the rate of internalization of the particles delivered to the cell. The rate of internalization, however, cannot exceed the rate of transport of particles to the cell (in other words, the rate of particle delivery), because the cell cannot internalize what is not there (causality). However, even if the rate of particle transport is high, the increase of cellular dose will be limited by the rate of internalization when particle internalization is sluggish. Cells that would internalize particles instantly would correspond to a Dirichlet boundary with vanishing concentration at the cells, and in this case the rate of internalization would be defined by particle transport.

Now we have our mathematical model completed, and next, we evaluate it and simulate expectable dosimetry functions.

### Evaluation of model

To compute our mathematical model and to simulate *in vitro* particle dosimetry curves in the geometry of 11-mm wells (Figure 2), we will use an initial concentration of *c*_0_ = 10 μg mL^−1^ (equal to 10 g m^−3^). Thus, with a filling-volume of 250 μL a total of 2.5 μg is administered to the well. The metric will be mass surface density (Supporting Information, Total dose), and we will also define and compute spatially-resolved dose (Supporting Information, Spatially-resolved dose). Spatially-resolved dose quantifies the dependence of the dose as a function of the distance from the center of the well. In this case, we divide the surface into 10 equal concentric parts (Figure SI 1b), and calculate the dose on each of the ten parts.

In the simulations we will use parameters corresponding to bare particles, although the surface of particles may be functionalized with synthetic polymers,^31^ or may become covered by proteins present in cell culture media,^32^ which may affect strongly the hydrodynamic radius and mass denisty.^6^ Nevertheless, from the point of view meniscus and its impact, the choice will simply boil down to the question whether diffusion or settling dominates the transport. The relative dominance of diffusion and settling velocity in the particle transport may be quantified by a dimensionless transport factor: 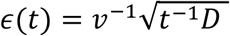. The amplitude of this factor is proportional to *∊ ∝ r_H_*^−2.5^Δ*ρ*^−1^, and *∊*(*t*) is a monotonically decreasing function of time. As long as *∊*(*t*) > 1, it is diffusion that dominates the transport, but when *∊*(*t*) < 1, the dominant way of transport is gravitational settling. In the limiting cases, *i.e.*, when either *∊*(*t*) ≫ 1 or *∊*(*t*) ≪ 1, the particles arrive to the cells only via diffusion and sedimentation, respectively. Accordingly, in the simulations we will use three different particle models (Small, Medium, Large) that correspond to typical systems and represent characteristic combinations of diffusion coefficients and settling velocities relevant for particle transport in quiescent fluids (Table 1).

**Table 1.** Particle models and their parameters used in this study to understand the impact of fluid menisci on *in vitro* dosimetry of particle dispersions.

| Particle model | Hydrodynamic diameter $2r_H$ & Mass density $\rho$ | Transport factor $\epsilon$ at $t = 1h$ |
| --- | --- | --- |
| **Small** (*e.g.*, Carbon dot) | 3 nm & 2 g/cm<sup>3</sup> | 34105 |
| **Medium** (*e.g.*, Gold particle) | 70 nm & 19 g/cm<sup>3</sup> | 0.72 |
| **Large** (*e.g.*, Polystyrene microsphere) | 1000 nm & 1.055 g/cm<sup>3</sup> | 0.31 |

For each particle model, we will use 14 bottom-boundary values *κ* (Neuman boundary condition, Equation 6). The three series of values of *κ* will be chosen by solving the three-dimensional mathematical model so that the total dose saturates within 1, 1.5…7 days up to the level of 99.5% of the administered dose. Figure 3a-c shows dosimetry curves obtained by *κ* values evenly spaced logarithmically over several orders of magnitude. Such values therefore mimic high, medium, and low affinities for particle internalization. Namely, saturation within one day mimics a high rate of particle internalization, where the increase of the dose is only limited by particle transport. Saturation within *e.g.*, seven days, however, mimics a low rate of particle internalization, where the dose-limiting process is not particle transport but particle internalization. Figure 3d shows the relationship found between the time of saturation and *κ* for the three particle models. We find that on a phenomenological level the relationships may be adequately described by a simple power-law model *a* + *b* ∙ *κ*^−*c*^, where the three model parameters may be obtained via nonlinear regression (Table 2).

**Table 2.** Particle models and the related best estimates of the model parameters used for characterizing the relationship between saturation time (99.5%) and the bottom-boundary value *κ* describing the affinity for particle internalization. In all the three cases, the quality of fit, quantified by the coefficient of determination (*R*^2^), is above 0.999.

| Particle model | Model parameter best estimates: $a + b \cdot \kappa^{-c}$ | | |
| --- | --- | --- | --- |
| | $a$ (day) | $b$ ($day\ m^{-c}$) | $c$ (1) |
| **Small** (e.g., Carbon dot) | $0.80 \pm 0.01$ | $(89.3 \pm 3.44) \times 10^{-9}$ | $1.02 \pm 0.00$ |
| **Medium** (e.g., Gold particle) | $0.73 \pm 0.02$ | $(8.09 \pm 2.62) \times 10^{-10}$ | $1.11 \pm 0.02$ |
| **Large** (e.g., Polystyrene microsphere) | $0.98 \pm 0.01$ | $(30.6 \pm 7.43) \times 10^{-12}$ | $1.15 \pm 0.01$ |

**Figure 3.**
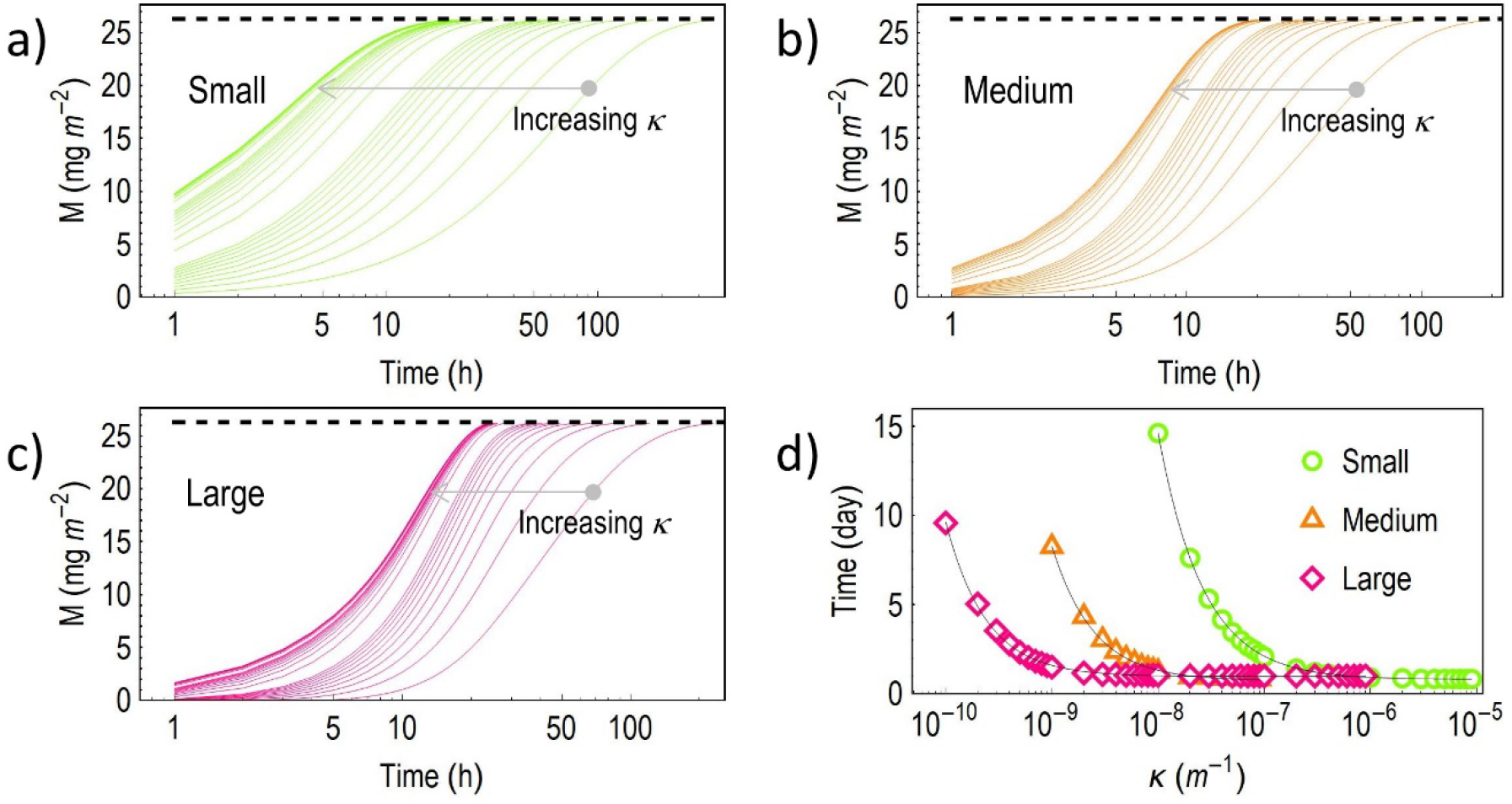
a)-c) Solutions of the three-dimensional model obtained at systematically varied bottom-boundary conditions. These values, spanning over several orders, represent a gradual shift from very low affinity to high affinity for particle internalization as the value of *κ* increases. The dashed lines indicate the administered dose. d) The dependence of the saturation time (99.5% of administered value) on the boundary value *κ*. The data points may be adequately described by power laws, and the solid lines are obtained via unconstrained nonlinear regressions by the method of least-squares providing the best model parameters estimates listed in Table 2.

Using these results, we solve the transport equation with the selected series of *κ* values that will result in the saturation of the dose within 1, 1.5…7 days. Figure 4-6 displays in detail selected but representative results obtained by our model with saturation in 1, 2 and 7 days.

**Figure 4.**
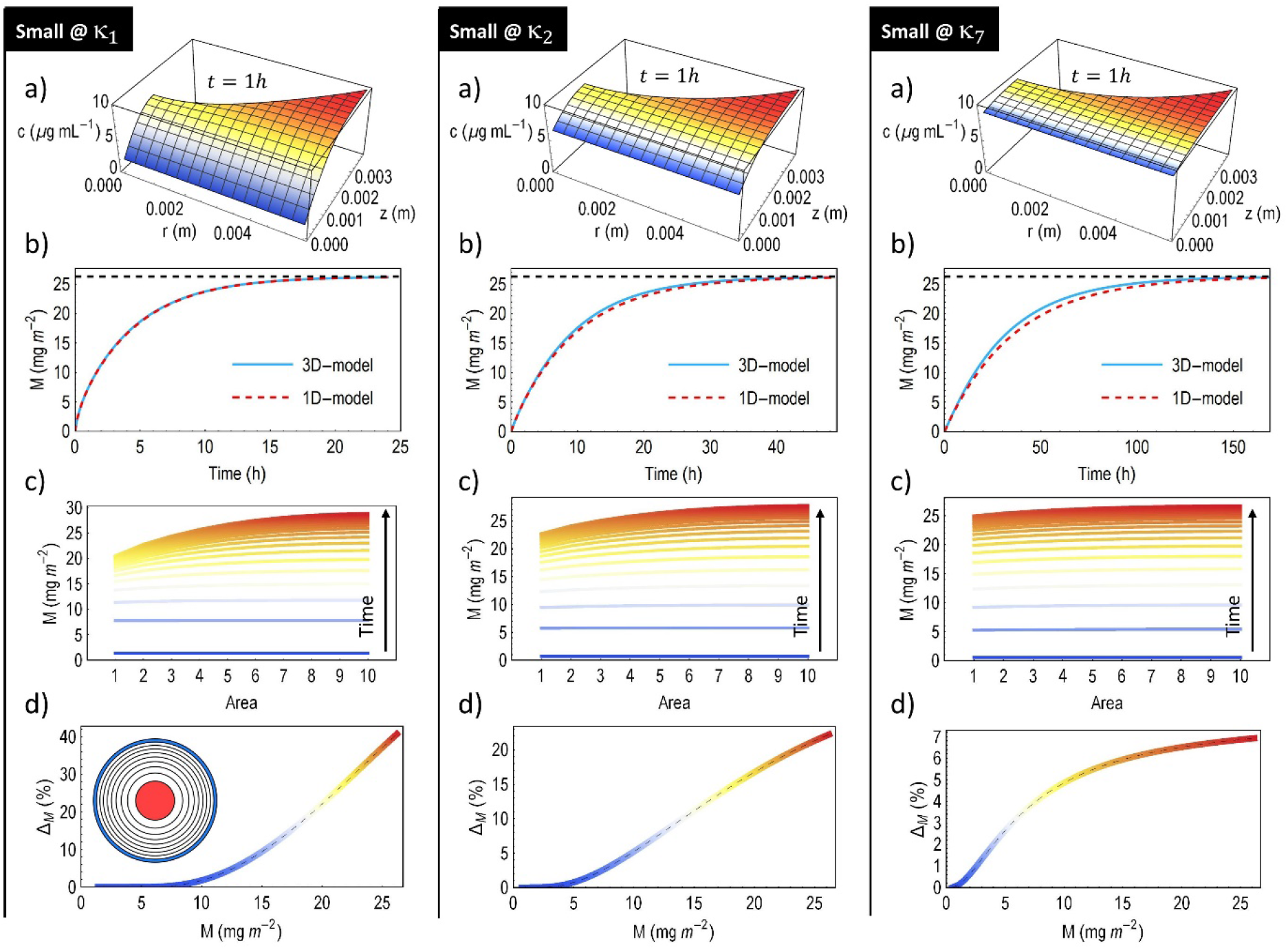
Representative results obtained by the ‘Small’ particle model. The value of the three internalization parameters: *κ* _1_, *κ* _2_, and *κ* _7_ are obtained via the mathematical model and the parameters shown in Table 2. These Neuman bottom-boundary conditions (Equation 6) result in saturation in 1, 2 and 7 days. Panels a) show snapshots of the concentration *c*(*t, r, z*) after one hour as a function of the cylindrical coordinates. Panels b) displays the total dose as a function of time (Supporting Information, Total dose) up to 1, 2 and 7 days. Panels c) show the evolution of spatially-resolved dose computed from the time- and space-dependence of *c*(*t, r, z*) on ten concentric parts of equal areas defined by increasing radius (Figure SI 1b, Supporting Information, Spatially-resolved dose). Panels d) show the maximum degree of nonuniformity— defined by the innermost (red) and the outermost (blue) parts each with a one-tenth of the total area—as a function of the total dose.

### Analysis of simulated results

The panels a) of Figure 4-6 show that particle concentration *c* is a nontrivial function of both time and spatial coordinates. The particle dose is calculated from *c*(*t, r, z*) as detailed in the Supporting Information file (Total dose, Spatially-resolved dose). Panels b) displays the total dose as a function of time up to 1, 2 and 7 days. Deviations of the result of the one-dimensional model from the three-dimensional model results from the fact that the solution of the one-dimensional transport equation does not scale linearly with height, and the one-dimensional model uses a volume-equivalent average fluid height (Supplementary Information, Nonlinear scaling of the solution of the one-dimensional transport equation). Nonetheless, these deviations are rather moderate. Panels c) show the evolution of spatially-resolved dose computed from the time- and space-dependence of *c*(*t, r, z*) on ten concentric parts of equal areas defined by increasing radius (Figure SI 1b). The dose is not uniform and increases systematically with distance from the center, which is particularly evident when the total dose is closing on saturation. Accordingly, cells grown near to center are expected to receive a dose smaller than cells grown near to the edge of the well do. As a matter of fact, the difference may be substantial and exceed 100% when particles settle heavily, *i.e.*, the outer cells may be expected to receive more than the double of the dose inner cells receive (Figure 5 and 6). The difference in dose is less critical in the case of particles whose transport is also influenced by translational diffusion (Figure 4). In such cases even a tendency can be identified: the longer the saturation takes, the less the nonuniformity. Indeed, Figure 7a shows that the degree of nonuniformity is a nontrivial function of the bottom-boundary. For the large particle model—where translational diffusion does not contribute to particle transport, and essentially gravitational settling is the only transport mechanism—the dependence is weak, and the maximum degree of nonuniformity Δ_*M*_(*t*) at saturation remains high independently on the affinity for particle internalization. However, when translational self-diffusion is not negligible in the transport of particles, the relationship between Δ_*M*_(*t*) at saturation and *κ* is strikingly different: the larger *κ*, the larger the nonuniformity. In other words, saturation that takes longer results in smaller dose nonuniformity (Figure 7b). This tendency may be understood by considering the role of diffusion in dispersing the administered particles across the surface parallel to orientation of cells, *i.e.*, in the direction defined by the radial coordinate *r*. When sedimentation dominates, transversal dispersion in any direction perpendicular to the pull of gravity is completely negligible, and particles simply descend onto the cell (Figure 8a). Consequently, in such cases, a higher fluid level above a given area means a greater mass of particles. This phenomenon was indeed observed by using fluorescence-labelled particles.^33^ For small particles, however, diffusion may effectively disperse particles before arriving to the cells (Figure 8b), especially if the affinity for particle internalization is small and the internalization is slow enough. In such cases, a) the longer it takes to saturate the dose the less the nonuniformity, and b) the higher the diffusion coefficient the less the nonuniformity (Figure 7b). Furthermore, given that different cell culture plates carry wells of a different dimension (Supplementary Information, Equilibrium shape of the meniscus, Table SI 1), we point out that the overall fluid volume may play a role in nonuniformity (Figure 8c). Finally, nonuniformity in dose systematically increases the variability of the responses of cells, which heterogeneity may exceed effect due to cell-to-cell variations within a population. Whether or not nonuniformity should be a matter of great concern will strongly depend on the overall dose, as illustrated in Figurer 8d. Dose-response relationship results in a measurable effect being a function of dose. Such function is generally nonlinear, and contain steeply and mildly increasing segments. Accordingly, the nonuniformity of the dose, *e.g.*, the difference between the dose near to the wall and near to the center of the well, may result in a considerable different cellular response and differing magnitude of the effect measured by analytical means.

**Figure 5.**
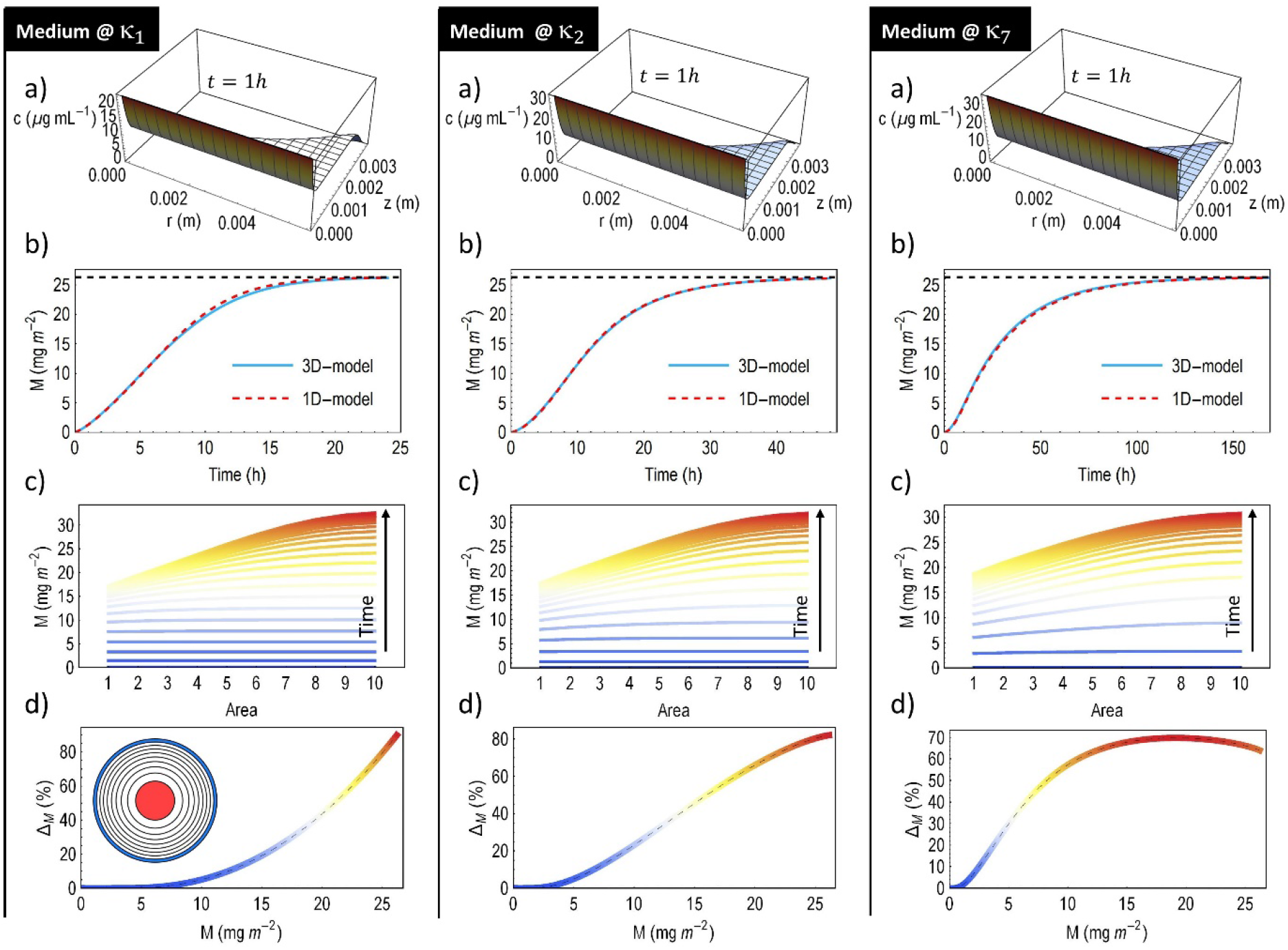
Representative results obtained by the ‘Medium’ particle model. The value of the three internalization parameters: *κ* _1_, *κ* _2_, and *κ* _7_ are obtained via the mathematical model and the parameters shown in Table 2. These Neuman bottom-boundary conditions (Equation 6) result in saturation in 1, 2 and 7 days. Panels a) show snapshots of the concentration *c*(*t, r, z*) after one hour as a function of the cylindrical coordinates. Panels b) displays the total dose as a function of time (Supporting Information, Total dose) up to 1, 2 and 7 days. Panels c) show the evolution of spatially-resolved dose computed from the time- and space-dependence of *c*(*t, r, z*) on ten concentric parts of equal areas defined by increasing radius (Figure SI 1b, Supporting Information, Spatially-resolved dose). Panels d) show the maximum degree of nonuniformity— defined by the innermost (red) and the outermost (blue) parts each with a one-tenth of the total area—as a function of the total dose.

**Figure 6.**
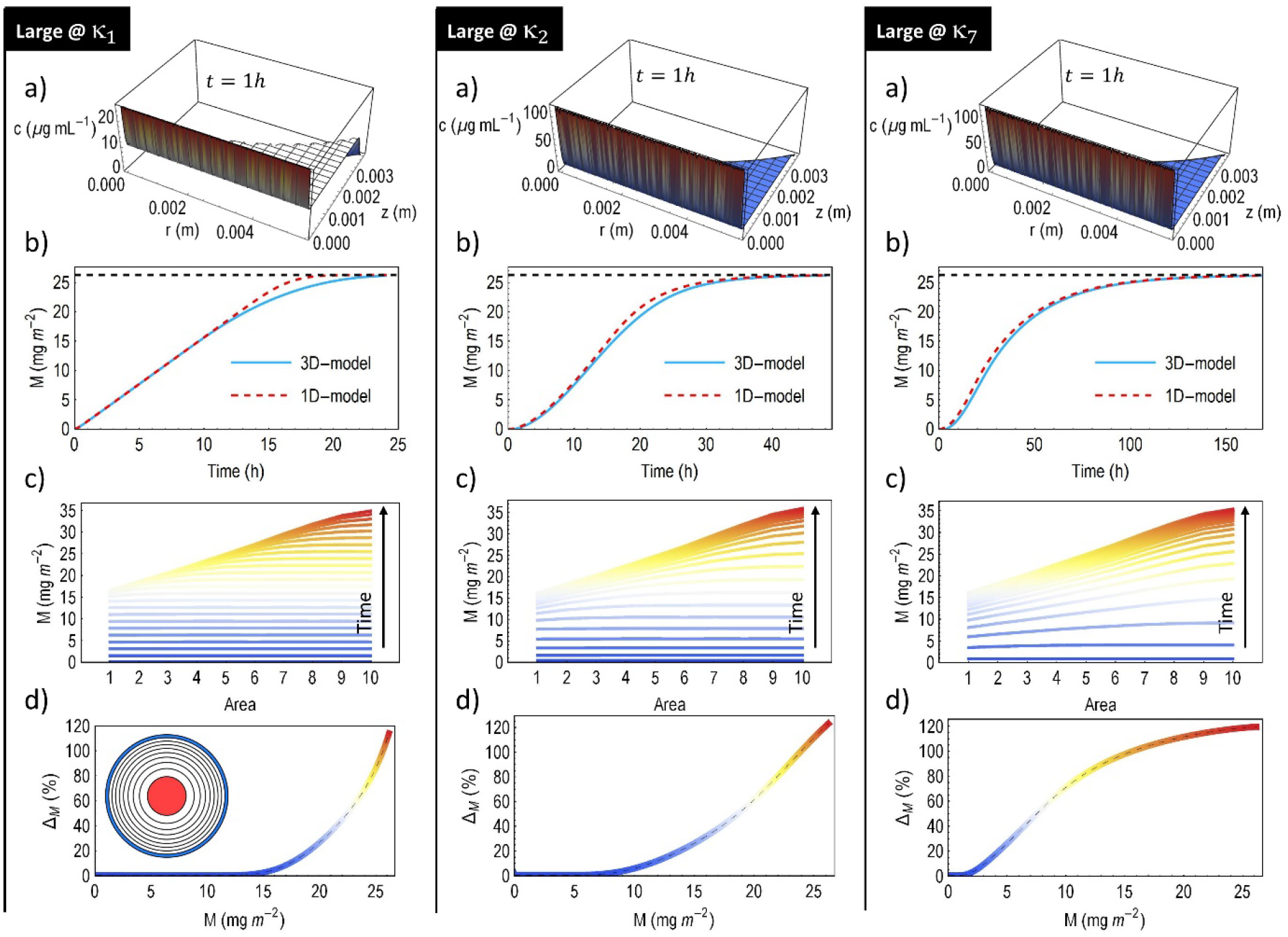
Representative results obtained by the ‘Large’ particle model. The value of the three internalization parameters: *κ*_1_, *κ*_2_, and *κ*_7_ are obtained via the mathematical model and the parameters shown in Table 2. These Neuman bottom-boundary conditions (Equation 6) result in saturation in 1, 2 and 7 days. Panels a) show snapshots of the concentration *c*(*t, r, z*) after one hour as a function of the cylindrical coordinates. Panels b) displays the total dose as a function of time (Supporting Information, Total dose) up to 1, 2 and 7 days. Panels c) show the evolution of spatially-resolved dose computed from the time- and space-dependence of *c*(*t, r, z*) on ten concentric parts of equal areas defined by increasing radius (Figure SI 1b, Supporting Information, Spatially-resolved dose). Panels d) show the maximum degree of nonuniformity—defined by the innermost (red) and the outermost (blue) parts each with a one-tenth of the total area—as a function of the total dose.

**Figure 7.**
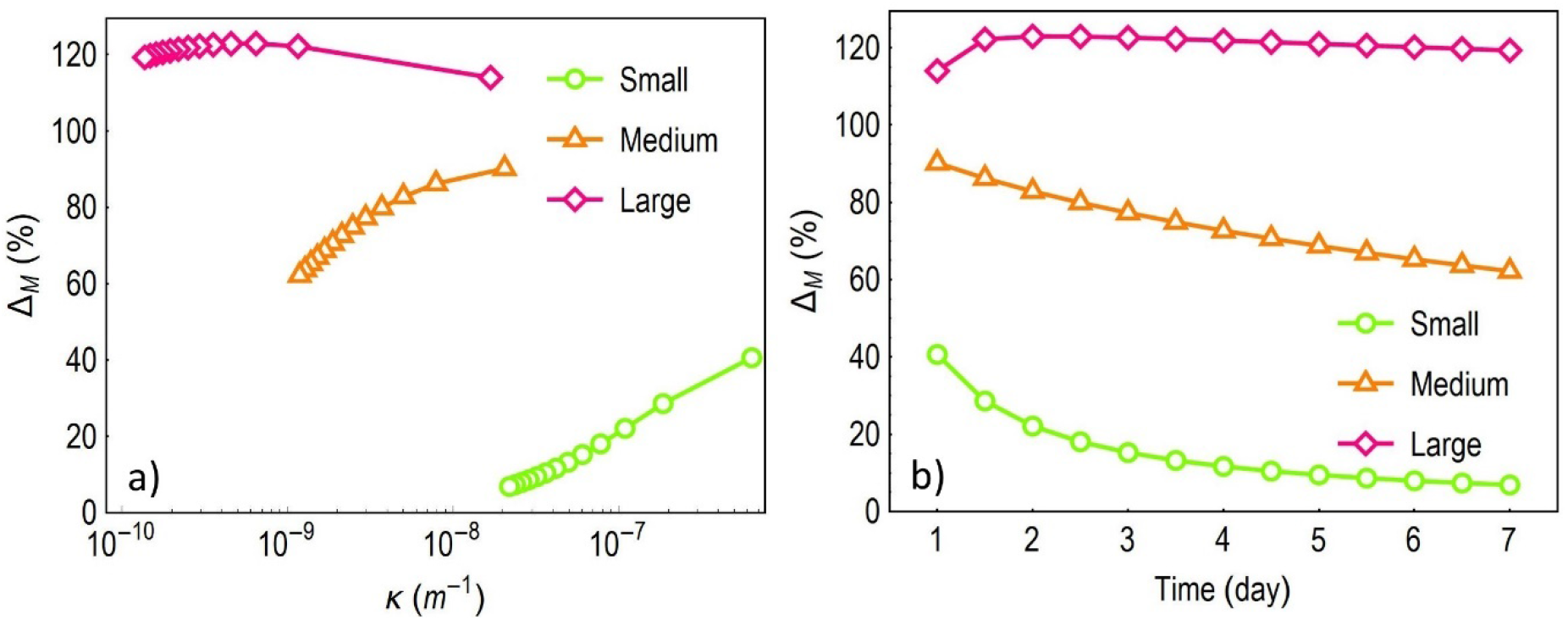
a) The maximum degree of nonuniformity at saturation as a function of the internalization parameter. b) The maximum degree of nonuniformity at saturation as a function of saturation time, computed at 1, 1.5…7 days.

**Figure 8.**
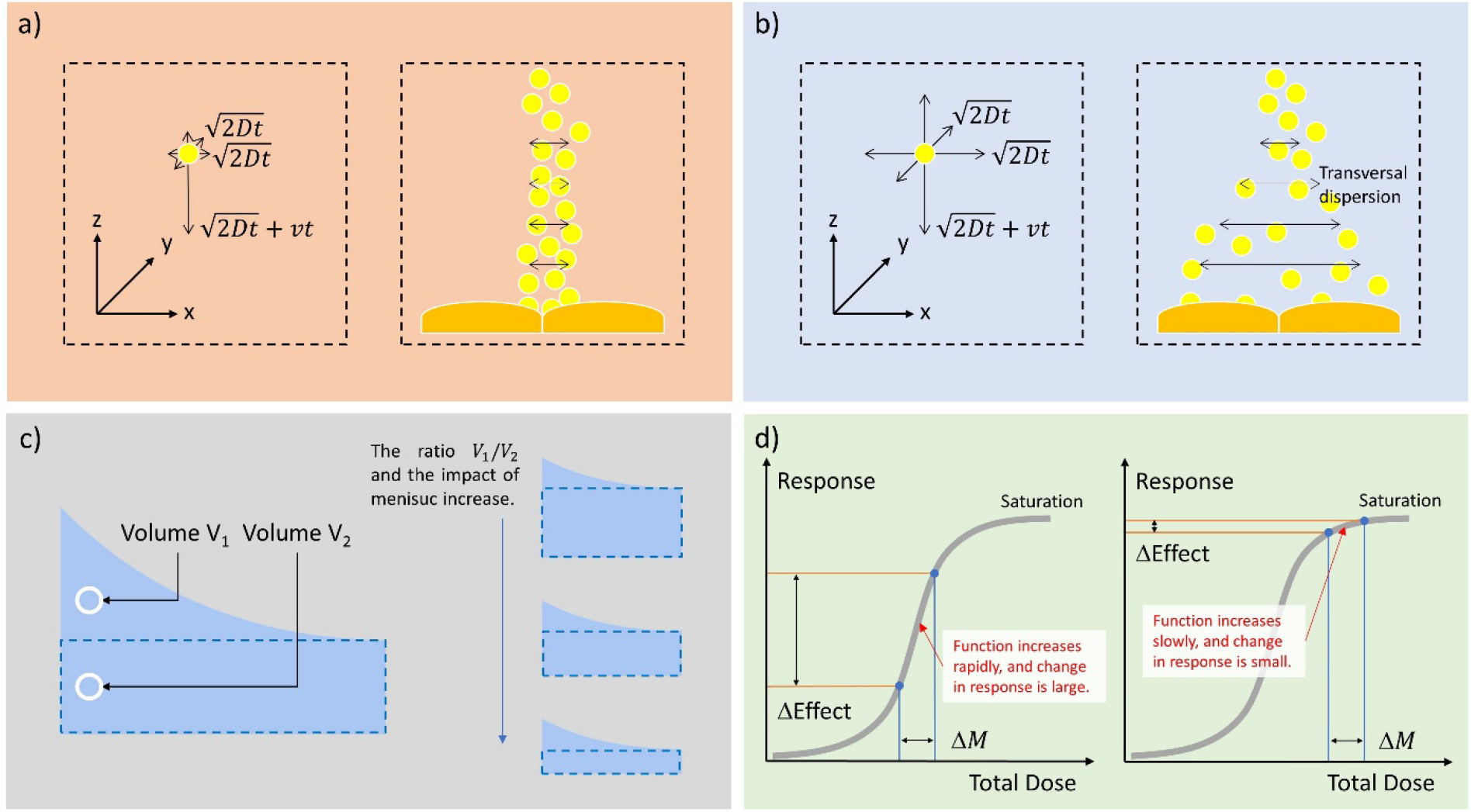
Processes and factors affecting the degree of spatially uneven dose, owing to fluid meniscus. a) Particles descend onto the cells without being dispersed transversally when sedimentation dominates particle transport. b) Particles may be dispersed transversally before arriving to the cells if diffusion is not negligible. c) The total fluid volume administered to a well may be divided into two sub-volumes: the volume defined by the curved meniscus (*V*_1_) and the volume under the meniscus (*V*_2_). Accordingly, the larger the *V*_1_/*V*_2_ ratio, the larger the nonuniformity. This ratio, however, decreases with overall fluid volume because *V*_1_ does not change as *V*_2_ increases—as long as the well is not full. d) Dose-response relationship results in a measurable effect being a function of dose. Such function is generally nonlinear, and contain steeply and mildly increasing segment over some *ΔM* interval before reaching saturation. Accordingly, a difference in the dose (Δ*M*), *e.g.*, difference between the dose near to the wall and near to the center of the well, may result in a considerable bias in the effect. In other words, the response of cells near to the wall and near to the center of the well may be considerably different if the total dose falls onto a steep region (left vs. right panel).

## SUMMARY

To summarize, fluid meniscus is a ubiquitous phenomenon in cell cultures, yet the impact on *in vitro* particle dosimetry has not been not addressed until now. We assembled a high-fidelity mathematical model, which was then exhaustively evaluated and analyzed. While no model is able to capture accurately all the details of each and every scenarios of *in vitro* particle exposures, we could identify and analyze the key factors that impacts the dose and its nonuniformity along the cell culture well. These findings also mean that the current state-of-the-art computational models are insufficient for they lack the ability to capture spatially uneven dosing, and consequently, they are not able to respect the fundamental concept of nanotoxicology. Accordingly, the next generation of *in vitro* particle dosimetry computational models must account for the meniscus.

## Acknowledgement

SB is grateful for the financial support of the Adolphe Merkle Foundation and the University of Fribourg, and also benefitted from support of the Swiss National Science Foundation through the National Center of Competence in Research Bio-Inspired Materials (25%). The authors thank Dr. Barbara Drasler for the original photograph used to prepare Figure 1.

