## Supporting Information for "Fluid Menisci and *In Vitro* Particle Dosimetry of Submerged Cells"

**Equilibrium shape of the meniscus**

The equilibrium shape of the meniscus may be obtained by solving the nonlinear second-order ordinary differential equation obtained via combining the principle of minimum energy with the Young–Laplace equation. A surface with axial symmetry, such as the meniscus with circular cross section in a cylindrical well, enables us to use cylindrical coordinates instead of Cartesian coordinates (Figure 2). Accordingly, owing to axial symmetry, the dependence on the angular coordinate drops out, and the problem becomes centrally symmetric and reduces to a one-dimensional nonlinear differential equation, where the height of the fluid surface is only a function of the distance from the center of the cylindrical well:[1]

(SI 1) $$f(r) = \frac{\gamma}{\rho g}\left(\frac{\partial_r^2 f(r)}{\sqrt{(1+(\partial_r f(r))^2)^3}} + \frac{\partial_r f(r)}{r\sqrt{1+(\partial_r f(r))^2}}\right).$$

Equation 7, however, does not have a straightforward analytical solution. Yet, given the dimensions of the wells commonly used in *in vitro* exposures, we may safely assume that the Gaussian curvature of the surface is small, and the square of the derivative is much smaller than one: $(\partial_r f(r))^2 \ll 1$. Then Equation 7 simplifies:

(SI 2) $$f(r) \cong \frac{\gamma}{\rho g}\left(\partial_r^2 f(r) + \frac{\partial_r f(r)}{r}\right).$$

To obtain the equilibrium shape of the meniscus, we solve Equation SI 2 along with the boundary conditions of $f(r=0) = z_0$ and $\partial_r f(r)|_{r=0} = 0$, where $f_0$ is the minimum fluid height in the center of the well, where it is locally flat with a zero Gaussian curvature. The solution is

(SI 3) $$f(r) \cong f_0 \cdot I_0\left(r\sqrt{\frac{\rho g}{\gamma}}\right)$$

where $I_n$ is the modified Bessel function of the first kind with $n = 0$, $\rho$ the mass density of the fluid, and $\gamma$ surface tension. Surface tension is a function of temperature, and in the case of an aqueous cell culture medium of 37 °C, $\gamma \cong 7 \cdot 10^{-4}$ J/m$^2$. For a given volume $V$ filling the well, we find the value of $f_0 = f(r = 0)$ via the following expression:

(SI 4) $$V = \int_0^R 2\pi \cdot r \cdot f(r)\, dr$$

where after performing the integration, we obtain

(SI 5) $$V \cong 2\pi \cdot R \cdot f_0 \sqrt{\frac{\gamma}{\rho g}} I_n\left(R\sqrt{\frac{\rho g}{\gamma}}\right)$$

where $I_n$ is the modified Bessel function of the first kind with $n = 1$. It is worth mentioning that for one-dimensional models, the volume-equivalent fluid height is $h = V/(\pi \cdot R^2)$, which is the ratio of the fluid volume and the surface area of the well. It is it is easy to show that $f(0) < h < f(R)$.

At last, the shape of the meniscus may vary slightly from well to well according to Equation SI-1, and below the dimensions and typical working volumes of the most commonly circular wells are listed.

**Table SI 1**. Dimensions and the working volumes of wells with axial symmetry and flat bottom.

| Number of wells | Shape | Diameter (mm) | Working volume (mL) |
|---|---|---|---|
| 6 | Cylindrical | 34.8 | 1.9-3 |
| 12 | Cylindrical | 22.1 | 0.75-1.5 |
| 24 | Cylindrical | 15.6 | 0.35-0.6 |
| 48 | Cylindrical | 11 | 0.2-0.5 |
| 96 | Truncated cone | 6.9 / 6.4 (top / bottom) | 0.15-0.2 |

### Transport equation

In quiescent and dilute dispersions inter-particle interactions are negligible, and the transport of nanoparticles is dependent on two quantities: translational diffusion coefficient $D$ and settling velocity $v$, which may be generally expressed as

(SI 6) $$D = k_B \cdot T/(\eta \cdot F)$$

(SI 7) $$v = g \cdot \Delta\rho \cdot V/(\eta \cdot F)$$

where $k_B$ is the Bolztman constant, $T$ temperature, $\eta$ viscosity of the fluid, $g$ gravitational acceleration constant, $\Delta\rho = \rho - \rho_f$ the difference in the mass density between the particle ($\rho$) and the fluid ($\rho_f$), $V$ volume of the particle.[2] When $\Delta\rho < 0$, the particles are buoyant, and do not settle but move upwards driven by buoyant force.[3] The term $\eta \cdot F$ is referred to as the friction coefficient, where $F$ itself is a function of particle dimensions and shape. Using accurate hydrodynamic models is necessary to successfully predict the hydrodynamic properties of anisotropic particles,[4] and the friction factor and effective mass density may be also obtained directly by experimental means.[5-17] In the case of spherical particles of hydrodynamic radius $r_h$, $F(R) = 6 \cdot \pi \cdot r_h$, and we obtain the well-known expression for the translational self-diffusion coefficient[18-20]

(SI 8) $$D = \frac{k_B T}{6 \pi \eta} \frac{1}{r_h}$$

and settling velocity

(SI 9) $$v = \frac{2g}{9\eta} \cdot \Delta\rho \cdot {r_h}^2.$$

The so-called transport equation, which is a second-order linear partial differential equation, describes particle concentration as a function of position and time[21]

(SI 10) $$\partial_t c(t, \boldsymbol{r}) + \mathrm{div}\,\left(\boldsymbol{v} \cdot c(t, \boldsymbol{r}) - D \cdot \mathrm{grad}\; c(t, \boldsymbol{r})\right) = 0$$

where in the general formulation $\boldsymbol{v}$ and $\boldsymbol{r}$ are vectors, and div and grad indicate are the well-known differential operators acting on vectors and scalars, resulting in scalar and vector quantities, respectively. Our metric is based on mass, and thus, $c$ represents mass concentration. The transport equation is used in a simple one-dimensional form, using the volume-equivalent fluid height (Figure 2)

$$\text{(SI 11)} \qquad \partial_t c(t,z) + v \cdot \text{div}\, c(t,z) - D \cdot \text{div grad}\, c(t,z) = 0$$

where div $c(t,z) = \partial_z c(t,z)$ and div grad $c(t,z) = \partial_z^2 c(t,z)$.

For the three-dimensional model, we use cylindrical coordinates, where $\boldsymbol{r} = [r, \theta, z]$, $\boldsymbol{v} = [v_r, v_\theta, v_z]$. Since $\partial_\theta c = 0$ (axial symmetry), $c(t, \boldsymbol{r})$ is only the function of time, distance from the center, and height from the bottom, *i.e.*, $c(t, \boldsymbol{r}) = [t, r, z]$. While the settling velocity is a vector quantity, in this geometry only the $z$-component is not zero: $\boldsymbol{v} = [0,0, v_z]$. Using these properties and cylindrical coordinates, we can express the transport equation as

$$\text{(SI 12)} \qquad \partial_t c + v\, \partial_z c - D\left(\frac{1}{r}\partial_r c + \partial_r^2 c + \partial_z^2 c\right) = 0.$$

**Initial condition and the impact of pipetting**

With the initial condition, we take that the concentration is uniform throughout the fluid volume at $t = 0$, and the dispersion is quiescent. This is a straightforward scenario that has been commonly used.[2,22-31] In other words: the impact of initial pipetting on particle transport is neglected. Exposure begins with administering particles to cell culture wells, which involves pipetting, and pipetting results in fluid convection that carries along the particles (advection). Advection may exhibit turbulent and chaotic behavior, and such advective streamlines may exhibit complex dynamic structures with vortices. The medium is therefore not quiescent for some time, and advection owing to pipetting is able to transport the particles at a higher rate, which is in contracts with the assumption that particle transport happens only via translational self-diffusion and gravitational settling. Nevertheless, advection will slow down and eventually halt owing to viscous dissipation, *i.e.*, the kinetic energy of the fluid is dissipated by viscous friction heating the fluid. When this happens, particle settling velocity and translation self-diffusion coefficient will dominate again the transport. The time it takes is a nontrivial function of the way of pipetting. The dynamics of the fluids may depend on several factors describing the technique of pipetting. Accordingly, we expect that it matters a) whether or not the tip of the pipette is immersed during sample release, b) depth and position of immersed orifice, *e.g.*, near to the wall or at center of the well, c) angle of tip compared to the well walls, d) duration of sample release, e) orifice size, f) speed of ejecting the sample jet, g) volume of the administrated fluid, h) volume of fluid already present in the well, and i) well diameter. This general scenario is arguably difficult to model by pure analytical means, and for a quantitative description, one needs to revert to numerical solutions addressing specific combinations of the factors above.

There is, however, an easier phenomenological approach to account for the impact of pipetting on particle transport: one may introduce a time-dependent dispersion coefficient $D_P$.[32-

[37] In this case, the diffusion term in the transport equation becomes: $D(t,r) = D_P(t) + D(r)$, and as time passes advection slows down as a result of dissipating the initial kinetic energy by fluid viscous friction, and thus, $D_P$ approaches zero. Given the numerous factors of pipetting, to obtain an adequate closed-form expression of $D_P$ is beyond trivial, but numerical approaches may provide quantitative estimates.

Another aspect is that particles reaching the cell surface are not internalized instantaneously, and owing to pipetting, the momenta of the particles are high, and the advection rate and related flow-induced shear stress maybe too high for adherence.[38] However, there are reports claiming that cellular uptake is not a monotonic function of shear stress, and in fact, shear stress may increase particle internalization.[39] Furthermore, particle internalization is dependent on a multitude of variables and involves a variety of endocytic mechanisms. There are pieces of evidence indicating that particle shape, volume, material elasticity, surface chemistry, spatial arrangement and surface density of ligands, cell type, cell cycle, and experimental methods may strongly influence particle internalization and cellular dose.[40-59]

**Total dose**

To measure the degree of internalization of particles, we use the mass depleted from the particle dispersion. The mass present in the well is calculated by integrating over the volume of the fluid

(SI 13) $\mu(t) = \int_V c(t, r, z)\, dv.$

In the case of cylindrical geometry this integral may be written as a double integral:

(SI 14) $\mu(t) = \int_0^R \left(2\pi \cdot r \cdot \int_0^{f(r)} c(t, r, z)\, dz\right) dr,$

and the mass internalized is

(SI 15) $m(t) = \mu(0) - \mu(t).$

The metric of dose we use is surface mass density, *i.e.*, the mass internalized by unit surface area, which in the case of total dose becomes:

(SI 16) $M(t) = \frac{m(t)}{R^2\pi}.$

where $R$ is the radius of the cell culture well. Equation SI 16 assumes that the coverage from cells is 100 %. From Equation SI 13-16, particle number, mass and surface area may be calculated directly. For the one-dimensional model, which uses a volume-equivalent fluid height ($h$), the integral given by Equation SI 13 and 14 is simpler:

(SI 17) $\mu(t) = R^2\pi \int_0^h c(t, z)\, dz,$

and then Equation SI 15 and 16 are used as above.

### Spatially-resolved dose

To compute the internalization of mass as a function of the distance from the center ($r$) of the cylindrical cell culture, we rely on the total dose defined earlier. Let $dm(t) = m(t + dt) - m(t) > 0$ denote the mass internalized in a short time-interval between $t$ and $t + dt$. According to the boundary condition we chose to use at the fluid-cell interface, internalization is proportional to the concentration (Equation 6).

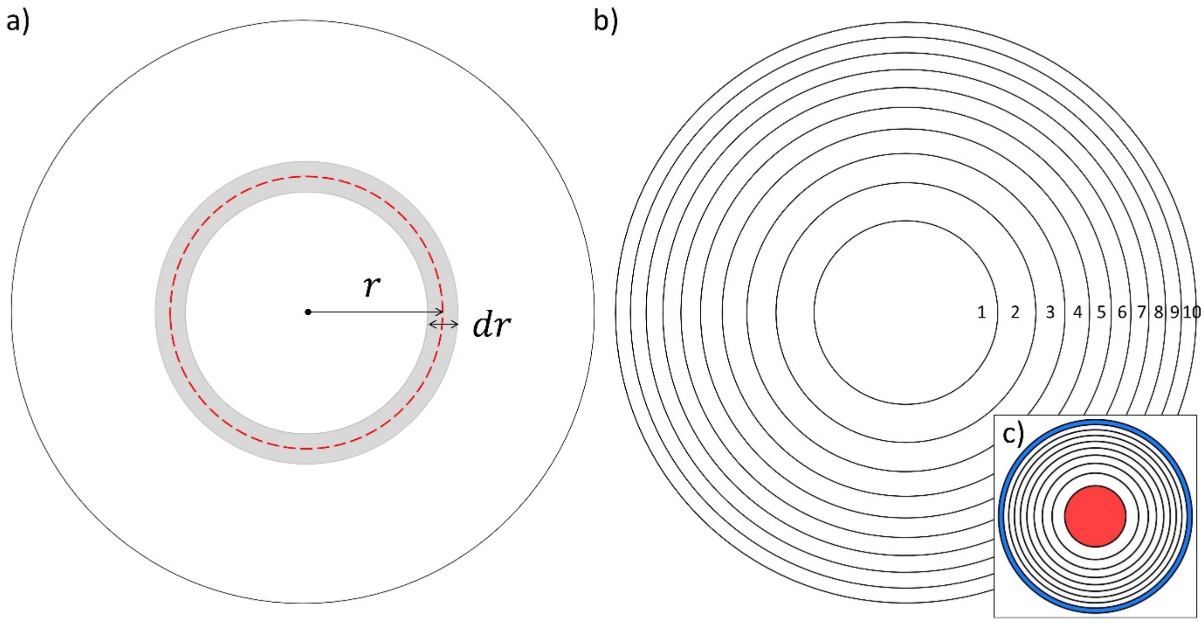

**Figure SI 1**. a) An annulus of radius $r$ and thickness $dr$. b) A central disk (1) and nine annuli (2-10) defining ten concentric parts of equal areas. c) The innermost (red) and the outermost (blue) parts each with a one-tenth of the total area.

The totality of particle mass internalized within $dt$ time is therefore obtained by integrating over the surface area of the cells: $\int_0^R dm(t,r)dr$, which may be written as

$$\text{(SI 18)} \qquad m(t + dt) - m(t) = \gamma \cdot \left( \int_0^R c(t,r,0) \cdot 2\pi \cdot r \cdot dr \right) dt.$$

Next, we can express $\gamma$ as

$$\text{(SI 19)} \qquad \gamma = \frac{m(t+dt) - m(t)}{dt} \left( \int_0^R c(t,r,0) \cdot 2\pi \cdot r \cdot dr \right)^{-1},$$

and by substituting Equation SI 20 into Equation SI 18, we obtain

(SI 20) $$dm(t,r) = \big(m(t+dt) - m(t)\big) \cdot \left(\int_0^R c(t,r,0) \cdot 2\pi \cdot r \cdot dr\right)^{-1} \cdot c(t,r,0) \cdot A(r).$$

The first and second term are to be evaluated from the solution of Equation 5 set up in the three-dimensional axially symmetric geometry (Figure 2), and $m(t)$ is given by Equation SI 15. From Equation SI 21 we can already see that the distribution of the mass internalized by the cells is not uniform, for there is a dependence on the radial coordinate, and thus, particle internalization is a function of the distance from the center of the well. Finally, we can express the mass internalized by an annulus of radius $r$ and area $A(r)$ until $t$ time

(SI 21) $$m(t,r) = \int_0^t dm(t',r) \cdot dt'.$$

Our metric is surface mass density, and thus

(SI 22) $$M(t,r) = \frac{1}{A(r)} \int_0^t dm(t',r) \cdot dt'.$$

Equation SI 24 defines the function describing the surface mass density as a function time (dose) of a narrow annulus of $r$ radius, such as the one depicted in Figure SI 1a. To compute spatially resolved dose on finite areas, we divide the surface into $n$ equal parts that are concentric (Figure SI 1b). Such an arrangement comprises one central disk and $n-1$ annuli, defined by a series of radii $r_i = R\sqrt{i/n}$, where $i = 1,2 \ldots n$. Accordingly, the area of any given part is $A_i = \pi({r_i}^2 - {r_{i-1}}^2) = \pi(R^2 i/n - R^2(i-1)/n) = \pi R^2/n$. In our case, we choose $n = 10$ (Figure SI 1b). At last, the maximum degree of heterogeneity in the spatial distribution of the dose is quantified via the innermost and outermost parts (Figure SI 1c):

(SI 23) $$\Delta_M(t) = \left(\frac{M(t,A_{10})}{M(t,A_1)} - 1\right).$$

**Nonlinear scaling of the solution of the one-dimensional transport equation**

The solution of the one-dimensional transport equation does not scale linearly with height, *i.e.*, if $c(z, t)$ is the solution for height $z = f(r)$, then $\lambda \cdot c(z, t) \neq c(\lambda \cdot z, t)$ where $\lambda$ is a positive real number, and $f(r)$ is the fluid height at coordinate $r$ (Equation 1, Equation SI 1). In other words, not only the amplitude, but also the shape of the dosimetry curve is dependent on the height. To show this, let us consider the dose from a volume defined by a narrow annulus of radius $r$ (Figure SI 1a) and height $f(r)$. The total mass contained initially in this cylindrical shell is $m_0(r) = c_0 \cdot f(r) \cdot 2\pi \cdot r \cdot dr$, where $c_0$ is the initial concentration. Let us consider instantaneous internalization and large particles (Table 1), where particle transport by diffusion is negligible ($\epsilon \ll 1$), and therefore, transversal diffusive transport (parallel to the surface of cells) and related diffusive mixing is vanishingly small. Accordingly, longitudinal transport (perpendicular to the surface of the cells) happens via sedimentation, and particle internalization is only limited by particle transport. In this case, the mass decreases linearly with time at a constant speed of settling, until the cylindrical shell empties out. Hence, the mass internalized as a function of time may be expressed as a piecewise function:

(SI 24) $$m_r(t) = \begin{cases} A \cdot t & t < \frac{f(r)}{v} \\ m_0(r) & t \geq \frac{f(r)}{v} \end{cases}$$

where $A = m_0(r)/f(r) \cdot v$, and the total dose is obtained by integrating the cylindrical shells:

(SI 25) $$m(t) = \int_0^R m_r(t).$$

The function expressed by Equation SI 27 exhibits a smooth saturation behavior, unlike the single one-dimensional function $m_r(t)$. Figure SI 2a shows three examples of $m_r(t)$, and Figure SI 2b shows the total dose.

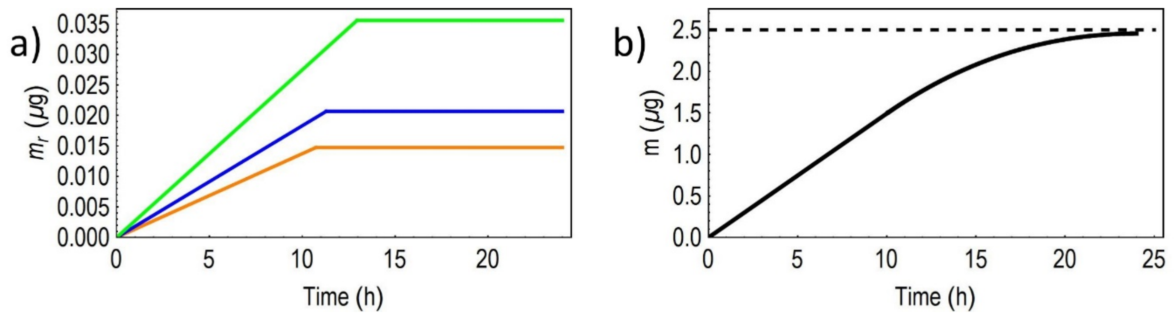

**Figure SI 2**. Dosimetry functions obtained by the solution of the one-dimensional transport equation with large particles and transport-limited particle internalization. The results are obtained with $r = R/4$ (orange), $R/3$ (blue), $R/2$ (green) and $dr = 10^{-4}$. b) The integral of the total dose was approximated by a finite sum of single functions.

(19) G., S. G. Collected Mathematical and Physical Papers, *Cambridge University Press,* **1880**, 75.
